# ProXs drive the formation of membraneless organelles in plants

**DOI:** 10.1101/2021.07.04.451080

**Authors:** Honghong Zhang, Fangyu Peng, Yan Liu, Haiteng Deng, Xiaofeng Fang

## Abstract

Membraneless organelles (MLOs) are non-membranous structures inside cells that organize cellular space and processes. The recent discovery that MLOs can be assembled via liquid-liquid phase separation (LLPS) advanced our understanding of these structures. However, the proteins that are capable of forming MLOs are largely unknown, especially in plants. In this study, we developed a method to identify proteins that we referred as ProXs (Proteins enriched by b-iso<u>X</u>) in Arabidopsis. Heterologous expression in yeast cells showed that most ProXs were capable of forming MLOs autonomously. We applied this method to several model and crop species including early and higher plants. This allowed us to generate an atlas of ProXs for studying plant MLOs. Analysis of ProXs from different species revealed high degree of conservation, supporting that they play important roles in cellular functions and are positively selected during evolution. Our method will be a valuable tool to characterize novel MLOs from desired cells and the data generated in present study will be instrumental for the plant research community to investigate MLO biology.

## Main

Eukaryotic cells are physically compartmentalized to ensure the efficiency and efficacy of biochemical reactions and cellular functions. Membraneless organelles (MLOs), which lack a surrounding lipid membrane, represent a unique way for compartmentalization. They confer a wide range of functional advantages over membrane-bound organelles due to their ability to spontaneously and rapidly form and dissolve. MLOs are super assemblies of proteins and frequently also nucleic acids that, in most cases, are assembled through phase separation^1,2^.

Within a eukaryotic cell, there are numerous MLOs that are both spatially and temporarily localized. However, only limited number of MLOs have been characterized. Some MLOs such as the nucleoli^3^, Cajal bodies^4^, P-bodies^5^, and stress granules^5^ are highly conserved and present ubiquitously in all eukaryotic cells. Some MLOs are specific to certain species or types of cells, performing highly specialized and essential functions. For example, germ granules only form in developing germ cells^6^ and the postsynaptic densities (PSDs) specifically assemble in postsynaptic neurons^7^. MLOs can have diverse sizes, compositions and material states determined by the collective behavior of their constituent macromolecules. However, for most MLOs, it has been proposed that usually only one or a few proteins, referred to as ‘scaffolds’, drive their formation. The other constituents, the so-called ‘client’ proteins, do not substantially contribute to the formation of the condensates^8^. Therefore, identification of scaffold proteins is key to characterizing the composition and function of novel MLOs.

Plants are quite unique due to their inability to move and need to rapidly cope with the continuous changes of light, temperature, water status, etc. This might have led to evolution of plant-specific MLOs that have yet to be discovered. Here, we developed a method to identify proteins that are prone to assemble MLOs *in vivo*. Using this method, we are able to characterize the MLO-forming proteins from the seedlings of eight plant species, ranging from early green alga to higher plants. Heterologous expression of these proteins confirmed that most of them indeed assemble MLOs on their own, therefore likely act as “scaffolds” for MLO formation. Our analysis revealed that the proteins responsible for MLO assembly are highly conserved during plant evolution. Our method and data provide new insights into cellular compartmentalization in eukaryotes, especially plants.

## Results

### Proteome-wide prediction of intrinsically disordered regions (IDRs) in Arabidopsis

Intrinsically disordered regions (IDRs) are protein segments that do not fold into a fixed three-dimensional structure yet exhibit biological activities^9^. Recent studies have linked MLOs with IDRs showing that the MLOs contain a significantly greater fraction of IDR-containing proteins (IDPs) than the overall proteome^10^. Accumulating evidence suggests that IDRs can be predicted from protein sequence features and dozens of computational methods based on different principles and computing techniques are available^11^. We selected two algorithms to predict IDPs in the proteome of Arabidopsis. IUPred is a physicochemical-based method that predicts disorder by assessing pairwise inter-residue interaction energy and identifies disordered residues as those that are incapable of forming stabilized inter-residue interactions^12,13^. The parameters of IUPred are derived from a global protein data set^14^, making it very robust and accurate. Two different sets of parameters were adopted to predict long (IUPred_L) and short (IUPred_S) disorder, respectively^13^. ESpritz is a machine-learning-based method that allows fast and accurate sequence-only prediction of IDRs^15^. It is trained by X-ray structure data set (ESpritz-Xray), NMR structure data set (ESpritz-NMR) or DisProt database (ESpritz-DisProt)^15^. The predictions showed that about 30%-45% of Arabidopsis proteins can be identified as IDPs (Extended Data Fig. 1a), similar to that of human proteome^16^. However, there is obvious discrepancy between different predictors, even when two predictors derived from the same algorithm (IUPred_L and IUPred_S) are compared (Extended Data Fig. 1b). These data suggest that it is difficult to define IDPs based on prediction and this could be more complicated *in vivo* because protein folding and unfolding is affected by numerous other factors. Therefore, it is not sufficient to identify MLO-forming proteins from predicted IDPs.

### Experimental method to identify MLO-forming proteins

We next sought to characterize proteins responsible for MLO formation by experimental approaches. The small molecule biotinylated isoxazole (b-isox) forms microcrystals that are able to precipitate proteins that localize in RNA granules^17^ as well as other MLOs^18,19^.

We therefore tested the ability of b-isox to enrich MLO-forming proteins in Arabidopsis. We incubated cell extract with various concentrations of b-isox at low temperature (4°C) and found that 100 μM b-isox precipitated proteins that were distinct from the total protein (Fig. 1a). This can be reversed by increasing the temperature to 37°C (Fig. 1a) that dissolves the b-isox crystals, indicating that the precipitation was mediated specifically by b-isox crystals. We then examined individual proteins that were known to assemble into MLOs *in vivo*, including FCA^20^, EMB1579^21^, SE^22^ and ELF3^23^, and found that they were all highly enriched by b-isox (Fig. 1b). In contrast, proteins that are unable to form MLOs were excluded from b-isox precipitates (Fig. 1b).

**Fig. 1.**
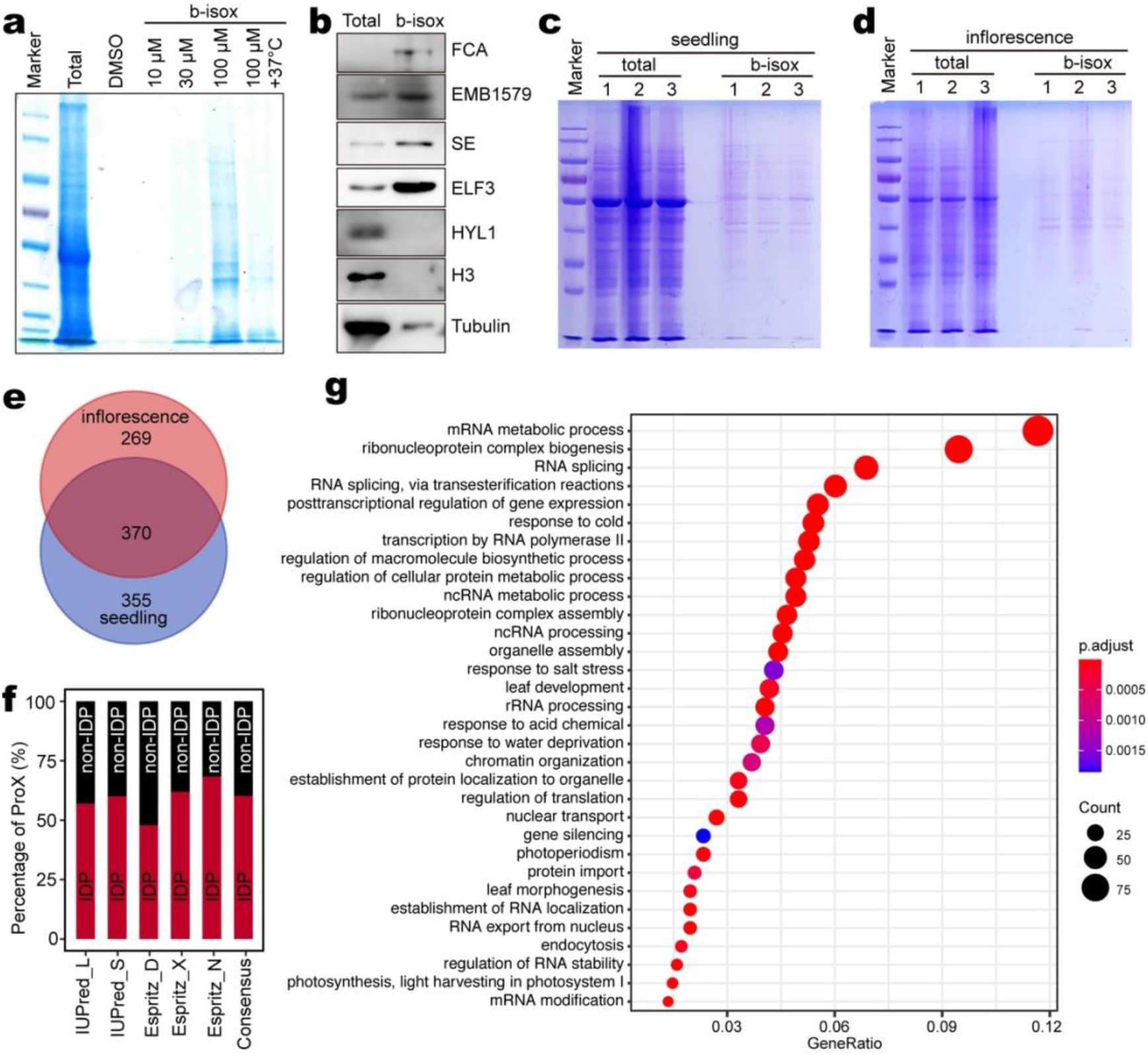
Identification of ProXs in Arabidopsis. **a**, Coomassie brilliant blue staining of an SDS-Polyacrylamide gel separating indicated protein samples. **b**, Detection of the indicated proteins in total and b-isox precipitation samples by western blot. **c**,**d**, Coomassie brilliant blue staining of SDS-Polyacrylamide gels separating total and b-isox precipitation samples prepared from Arabidopsis seedlings (**c**) and Arabidopsis inflorescences (**d**). The numbers indicate three biological replicates. **e**, Venn diagram showing the overlap of the ProXs identified from Arabidopsis seedlings and inflorescences. **f**, The percentage of IDPs contained in the ProXs shown in (**e**) as predicted by indicated programs. The consensus IDPs were defined by at least two programs. **g**, GO term analysis of the ProXs shown in (**e**).

The above results support the feasibility to characterize MLO-forming proteins using b-isox precipitation. Given that some proteins are expressed only at specific developmental stages, we separately prepared cell extracts from Arabidopsis seedlings and inflorescences and performed the b-isox precipitations (see Methods). Three biological replicates of both total protein and b-isox precipitates were subjected to mass spectrometry analyses (Fig. 1c, 1d). High reproducibility was observed between the replicates (Extended Data Fig. 2). One of the replicates from inflorescences b-isox precipitates, which showed poor correlation with two other replicates (Extended Data Fig. 2b), was excluded from subsequent analyses. After subtracting the background resulted from high abundance in total extract and filtering out the unconfident counts, a total of 725 and 639 proteins were significantly enriched by b-isox from seedlings and inflorescences, respectively (Fig. 1e, Supplementary Table 1). These proteins are referred to as ProX (<u>Pro</u>teins precipitated by b-iso<u>X</u>). Over half of the ProXs from seedlings and inflorescences overlapped with each other (Fig. 1e). We merged those overlapped ProXs, giving rise to 994 ProXs for Arabidopsis (*At*ProXs) (Supplementary Table 1). Among the ProXs, over 60% of them harbor IDR (Fig. 1f), which is significantly higher than the ratio in the proteome (Extended Data Fig. 1a). This supports a role of IDR in attaining the ability to be precipitated by b-isox^17^. Nucleolus, stress granules (SGs) and P bodies (PBs) are conserved MLOs that were also well-characterized in Arabidopsis^24–26^. Importantly, our ProXs included most of the proteins that are essential for the assembly of nucleolus, SGs and PBs (Supplementary Table 2). Moreover, we also identified other proteins that were recently reported to form MLOs (Supplementary Table 2). For example, FLOE1 formed cytoplasmic MLOs when expressed^27^ and GBPL3 formed GBPL defence-activated condensates (GDACs) that regulated plant immunity^28^. These data strongly suggest that our method indeed significantly identified the proteins involved in MLO formation.

The Gene Ontology enrichment analysis showed that most of the *At*ProXs were involved in RNA-related processes, including transcription, splicing and other post-transcriptional events (Fig.1g), consistent with a broad role of MLOs in RNA biology^29^. In addition, many *At*ProXs were enriched in responses to various stimuli, suggesting that MLOs may play key roles in stress responses. Indeed, there are numerous examples in which MLOs provide adaptation to specific environmental cues including temperature, pH, osmotic and nutrient changes^30^.

### Validation of ProXs as proteins scaffolding MLO assembly

It was not clear to what extent the proteins enriched by b-isox were able to scaffold the assembly of MLOs^17^. To test this, we took the advantage of a heterologous expression system in fission yeast cells, which stably integrates a gene of interest into an endogenous locus (Fig. 2a)^31^. We reasoned that because of the huge evolutionary distance between plants and yeast, the ability of a plant ProX to form MLOs in yeast cells, to a large extent, indicates that it acts as a scaffold rather than a client for MLO formation. RNA granules are ubiquitous MLOs that play essential roles in the regulation of gene expression at both transcriptional and posttranscriptional levels^32^. The major components of RNA granules are RNA-binding proteins (RBPs) and RNAs. In Arabidopsis seedlings, a total of 822 proteins have been identified as RBPs^33,34^, among which 284 of them were covered by our *At*ProX list (Fig. 2b, Supplementary Table 3). We fused these RBPs with green fluorescence protein (GFP) and expressed them in yeast cells. Out of 80 RBPs randomly chosen from our *At*ProX list (Supplementary Table 4), 13 of them did not give rise to detectable fluorescence signal possibly due to incompatible codon usage (Supplementary Table 4). Surprisingly, 94% (63 out of 67) of the expressed RBPs formed distinct foci (Fig. 2c, Extended Data Fig. 3). As a control, we also expressed 10 RBPs that were excluded by b-isox (Supplementary Table 5) and found that only one of them formed obvious foci (Fig. 2d, Extended Data Fig. 4). Taken together, these data support that the ProXs we identified represent very confident candidates that scaffold the assembly of MLOs.

**Fig. 2.**
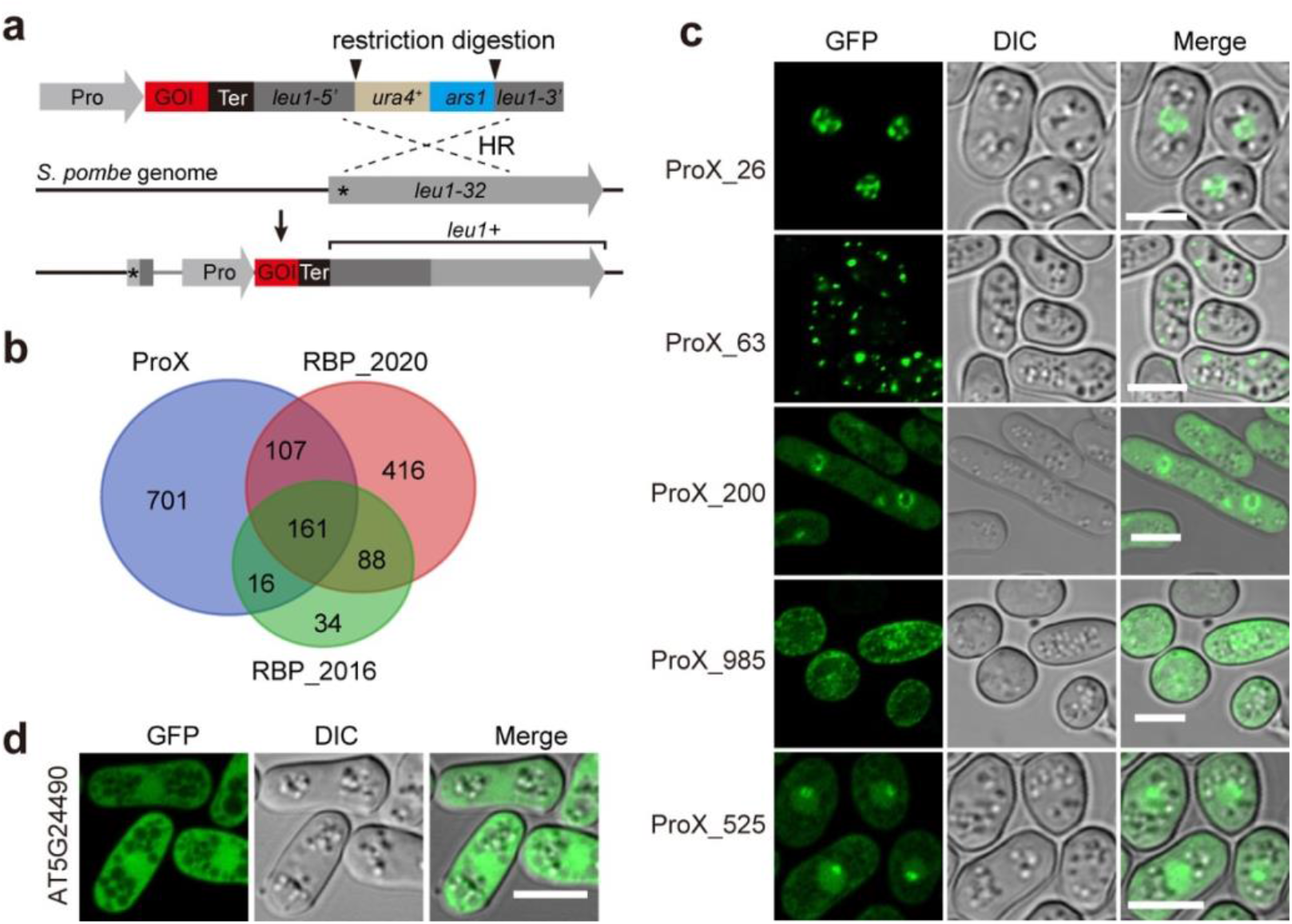
Validation of the ability of ProXs to form membraneless organelles (MLOs). **a**, Illustration of the yeast expression system (see Methods). The asterisk indicates the position of the *leu1-32* mutation. HR, homologous recombination. **b**, Venn diagram showing the overlap of Arabidopsis ProXs with RBPs identified by two independent studies. **c**, Representative images of ProXs that formed distinct foci in yeast cells. Scale bars, 5 μm. **d**, Expression of an RBP that was excluded by b-isox precipitation in yeast cells. formed distinct foci in yeast cells. Scale bar, 5 μm.

### Characterization of ProXs from other plant species

The effectiveness of our method to identify MLO-scaffold proteins prompted us to apply it to other plant species including the green alga *Chlamydomonas reinhardtii* (*Cr*), moss *Physcomitrella patens* (*Pp*), rice *Oryza sativa* (*Os*), maize *Zea mays* (*Zm*), common wheat *Triticum aestivum* (*Ta*), Chinese cabbage *Brassica rapa ssp. pekinensis* (*Br*) and tomato *Solanum lycopersicum* (*Sl*) (Fig. 3). We selected these species for two reasons: first, they are all crop and model plants important to the research community; second, they span a long evolutionary history, for example, *Chlamydomonas reinhardtii* is the unicellular green plant and *Physcomitrella patens* represents the earliest land plant. We collected materials at developmental stages that were similar to Arabidopsis seedlings and subjected to b-isox precipitation (Extended Data Fig. 5). The same procedure was applied to analyze all data (see Methods). Variable number of ProXs were identified, with *P. patens* having the most (1168 ProXs) while *C. reinhardtii* and *S. lycopersicum* having fewer (494 and 516 ProXs, respectively) (Fig. 3a, Supplementary Table 1). We also performed proteome-wide prediction of IDPs for all species. The overall ratio of IDPs in all species except *C. reinhardtii* is similar to that in Arabidopsis (Fig. 3b). *C. reinhardtii* contained exceptionally high ratio of predicted IDPs, with 70% of its proteome being IDPs (Fig.3b). Nevertheless, ProXs from all species contained higher ratio of IDPs compared to that in proteomes (Fig. 3c).

**Fig. 3.**
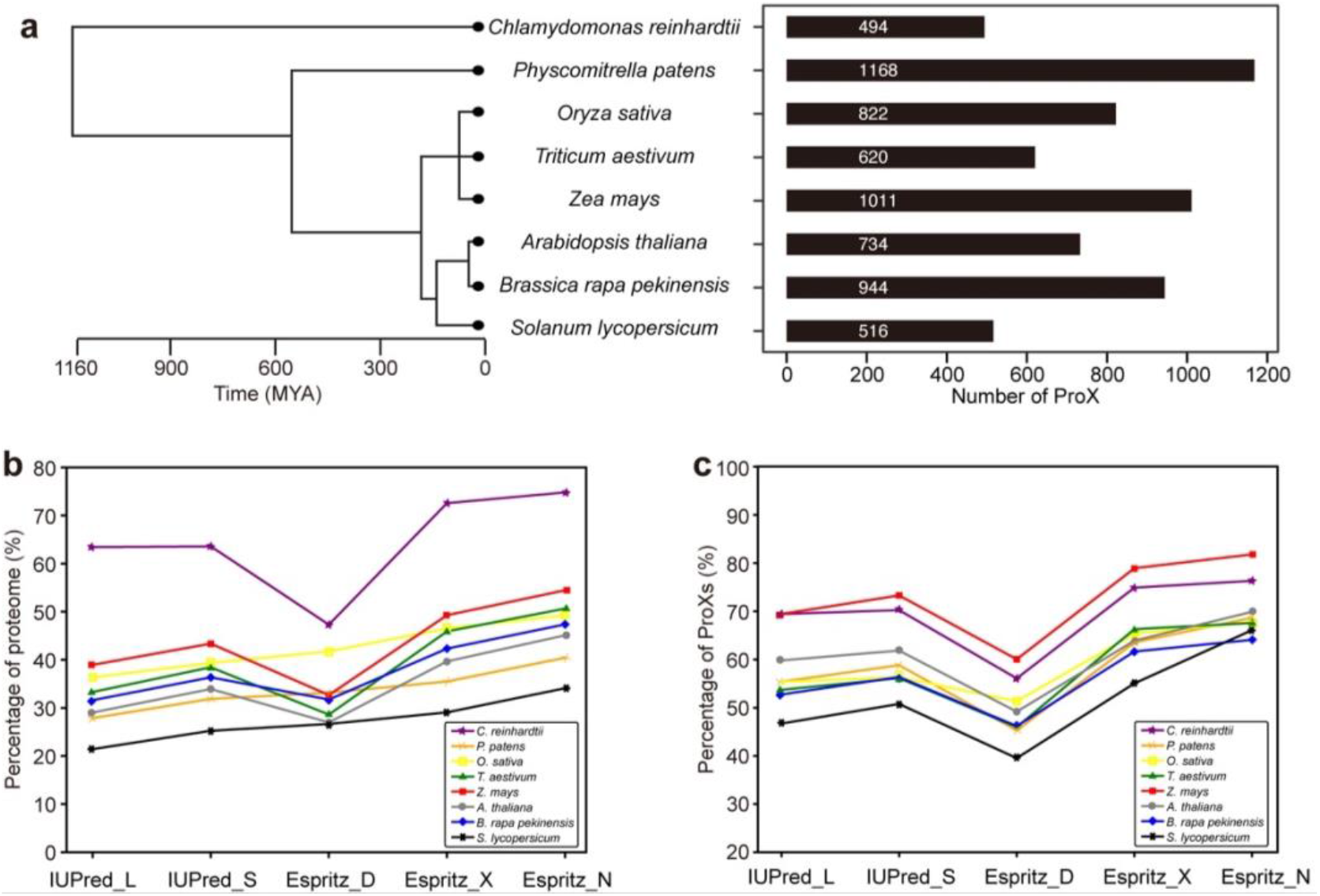
Characterization of ProXs from representative plant species. **a**, The phylogenetic tree of selected plant species (left) and the number of ProXs identified from each species (right). MYA, million years ago. **b**, The percentages of IDPs in the proteomes of indicated species as predicted by individual programs. **c**, The percentages of IDPs in the ProXs of indicated species as predicted by individual programs.

### Conservation analyses of ProXs

Given that the species we chose span a long evolutionary distance, it is worthy to perform a comparative analysis of the ProXs from these species. For comparison, we included the ProXs of Arabidopsis seedlings. We first assessed the functions of ProXs by GO enrichment analysis (Fig. 4a). The results showed that ProXs from all species were invariably enriched in RNA-related processes including transcription, RNA processing, splicing, etc. Interestingly, the terms such as endocytosis and photosynthesis were overrepresented in almost all species, suggesting a conserved role of MLOs during these processes.

**Fig. 4.**
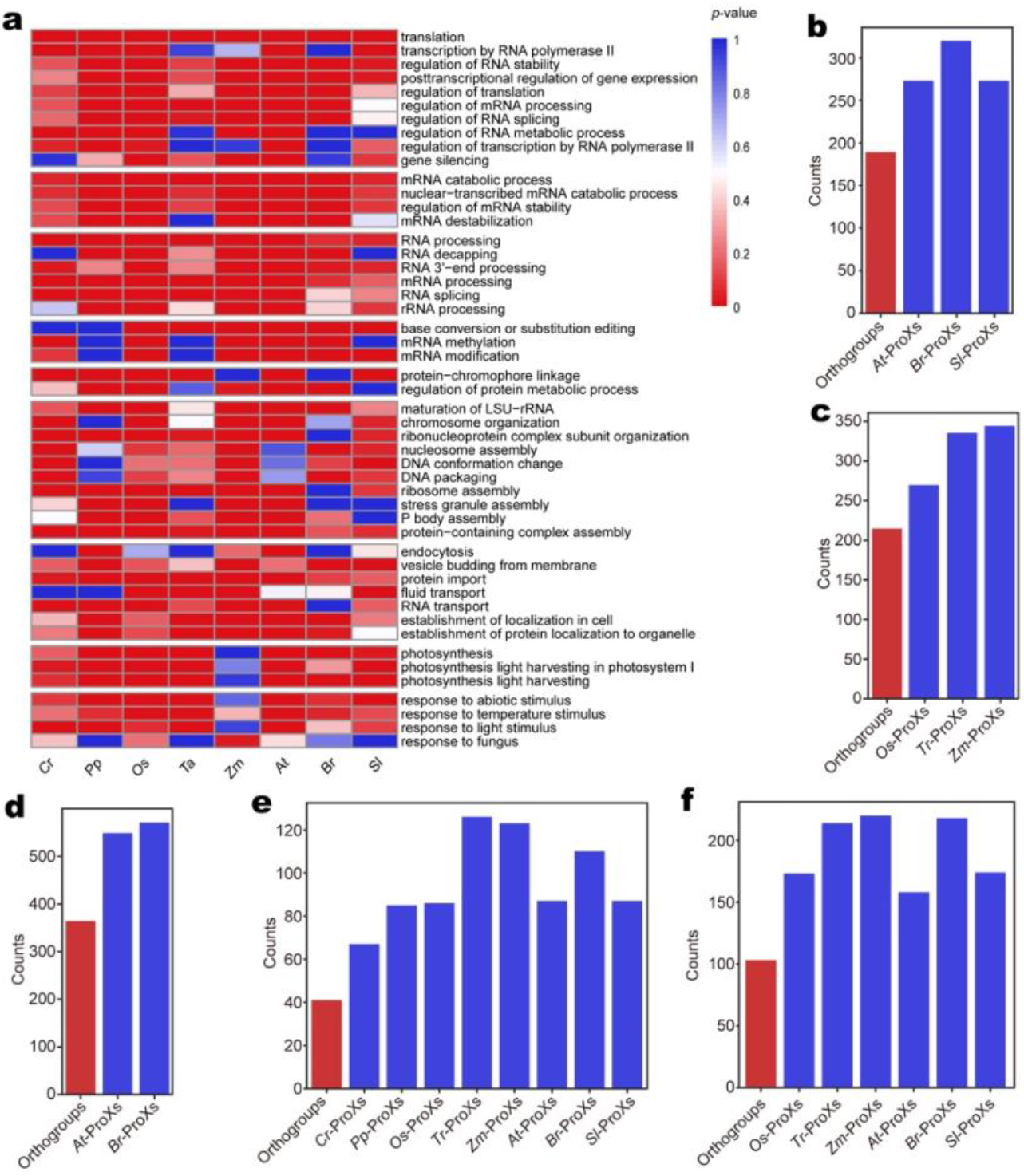
Analyses of ProXs from representative plant species. **a**, GO term analysis of the ProXs identified from eight species. **b**, The number of orthogroups identified within eudicots and the numbers of ProXs included by those orthogroups. **c**, The number of orthogroups identified within monocots and the numbers of ProXs included by those orthogroups. **d**, The number of orthogroups identified between *A. thaliana* and *B. rapa ssp. pekinensis* and the numbers of ProXs included by those orthogroups. **e**, The number of orthogroups identified between eight selected species and the numbers of ProXs included by those orthogroups. **f**, The number of orthogroups identified within angiosperm and the numbers of ProXs included by those orthogroups.

We then determined the phylogenetic relationships between ProXs from different species. We first analyzed the homology within eudicots and monocots, respectively. OrthoFinder (see Methods) defined 189 and 215 orthogroups inside eudicots and monocots, respectively (Fig. 4b,c, Supplementary Table 6). These orthogroups covered 1/3~1/2 of their ProXs. Within the same family Brassicaceae, *A. thaliana* and *B. rapa pekinensis* had 364 orthogroups, accounting for 549 (75%) and 571 (60%) of their ProXs (Fig. 4d, Supplementary Table 6). Fourty-one orthogroups were found when all eight species were compared (Fig. 4e, Supplementary Table 5). This number was relatively small mainly because nearly 40% of the *C. reinhardtii* ProXs had no homologs in the proteomes of other seven species therefore cannot be assigned to any orthogroup (Extended Data Fig. 6). Indeed, this number was increased to 103 when only higher plants were compared (Fig. 4f, Supplementary Table 6). The degree of homology between ProXs is underrepresented if we take into account that homologous ProXs may not be expressed at the developmental stage we used and therefore cannot be identified by b-isox precipitation. Intriguingly, multiple sequence alignment of ProXs within orthogroups showed that the sequence conservation between IDRs was very poor (Extended Data Fig. 7), consistent with the observation that the disordered regions evolved more rapidly than the ordered region^35^. These data suggest that MLO-forming proteins are positively selected during evolution. In addition, the above results support that our method is robust.

The research on MLOs caught increasing attention in the past decade thanks to the discovery that most MLOs assemble via LLPS. However, the study of MLOs in plants is still in its infancy. The ProXs we identified will serve as candidates to characterize novel MLOs and expand our understanding of the molecular mechanism and functional importance of LLPS. Since the composition of b-isox precipitates relies on the constituents of the input sample, our method is more advantageous to identified MLOs exist in specific tissues, cell types, or under certain conditions. For instance, ProXs enriched specifically under stress treatments are more likely to assemble stress-responsive MLOs and function in stress responses. In all, the data and method generated in present study will be valuable for the plant research community to investigate MLO biology.

## Materials and Methods

### Plant materials and growth conditions

For *Arabidopsis thaliana*, 12-day-old Col-0 seedlings grown on standard half-strength Murashige and Skoog (MS) (0.22% MS, 1% sucrose, 0.4% Phytagel, Sigma, P8169) media plates and whole flower buds from one-month-old Col-0 plants grown in soil were used. All plants were grown under long photoperiod conditions (16 h light of 120 μmol m^-2^ s^-1^/8 h dark). For *Brassica rapa subsp. pekinensis*, the 20-day-old seedlings of Chinese cabbage “Baiyang” were kindly provided by Dr. Tongbing Su. For *Solanum lycopersicum*, the 15-day-old seedlings of M82 ecotype were gifted by Dr. Cao Xu. For *Triticum aestivum and Zea mays*, 10-day-old seedlings of Chinese Spring and B73, respectively, were kindly provided by Dr. Zhaorong Hu and Dr. Feng Tian. For *Oryza sativa*, 14-day-old seedlings of Nipponbare ecotype were kindly provided by Prof. Yijun Qi. For *Physcomitrella patens*, the 7-day-old gametophytes of Gransden ecotype were kindly provided by Dr. Haodong Chen. For *Chlamydomonas reinhardtii*, cells of CC-1690 wild type mt+ strain grown to logarithmic phase were gifted by Prof. Junmin Pan.

### Proteome data collection and filtering

Protein sequences for the following eight plant proteomes were collected from UniProt (https://www.uniprot.org/): *Chlamydomonas reinhardtii* (*Cr*), *Physcomitrella patens* (*Pp*), *Oryza sativa* (*Os*), *Triticum aestivum* (*Ta*), *Zea mays* (*Zm*), *Arabidopsis thaliana* (*At*), *Brassica rapa pekinensis* (*Br*), and *Solanum lycopersicum* (*Sl*). Because the invalid residues (B, J, O, U, X, Z) cannot be recognizable by disorder prediction algorithms, proteins containing these amino acids were filtered out from our datasets. A summary of the proteome data of each species are shown in Supplementary Table 7.

### IDR prediction

Prediction of intrinsically disordered regions was performed using 5 predictors, including three versions of ESpritz (http://old.protein.bio.unipd.it/download/): ESpritz-Xray, ESpritz-NMR, ESpritz-DisProt and two versions of IUPred (http://iupred.enzim.hu/Downloads.php): IUPred-long, IUPred-short.

For each predictor, we first defined the disorder probability ranging from 0 to1 for each residue. All scores were then converted into a binary value to define a disordered residue according to the default threshold (IUPred-Long: 0.5, IUPred-short: 0.5, ESpritz-DisProt: 0.5072, ESpritz-Xray: 0.1434, and ESpritz-NMR: 0.3089). Regions with 30 or more consecutive disordered residues were classified as IDRs.

### Gene Ontology annotation and analysis

The gene ontology (GO) annotation of eight proteomes was retrieved from the UniProt-GOA database (https://www.ebi.ac.uk/QuickGO/). GO enrichment analysis was performed using the g:Profiler^36^—a web server for functional enrichment analysis. The options of significance threshold choose the Benjamini-Hochberg FDR, other options with default setting Bioconductor package clusterProfiler^37^ using the default parameters.

### Evolutionary analyses of ProXs

The amino acid sequences of ProXs were retrieved from proteomes of interest. The Orthofinder2.3.12^38^ was used to identify orthogroups between the species compared. The running script is *orthofinder -f datasets -M msa -S diamond -A muscle -t 8 -a 8*.

### Protein extraction and b-isox precipitation

All plant materials were ground in liquid nitrogen into fine powder. Total proteins were extracted using extraction buffer (10 mM Tris-Cl, 150 mM NaCl, 5 mM MgCl_2_, 20 mM β-Mercaptoethanol, 0.5% NP-40; 10% Glycerol, 1% protease inhibitor cocktail, pH 7.5). After rotation for 30 min at 4°C, the cell lysates were cleared by centrifugation (4°C, 14000 rpm for 15 min, twice) and the extract was filtered through 0.45 μm filter membranes (Millipore). The biotinylated isoxazole was synthesized as described previously^17^. Biotinylated isoxazole (B-isox)-mediated precipitation was performed according to Kato *et al*^17^. Briefly, a final concentration of 100 μM b-isox was add to 1.5 mL extract with protein concentration adjusted to 5 mg/mL. The extract was then incubated at 4°C for 1 h and centrifuged at 14000rpm for 15 min. The pellets were washed three times with ice-cold extraction buffer and boiled in 20 μL protein loading buffer at 95°C for 10 min.

### Immunoblot analysis

Protein samples were separated by SDS-PAGE gel and transferred to PVDF membranes. Antibodies against FCA^39^, SE (Agrisera, AS09532A), HYL1^40^, ELF3^41^, H3 (Abcam, ab1791), GFP (Roche, 11814460001), Tubulin (Sigma, T5168) were used as primary antibodies. After the primary antibody incubation, horseradish peroxidase (HRP)-conjugated secondary antibodies (GE Healthcare) were used for protein detection by chemiluminescence (Thermo Scientific, 34095).

### Yeast expression

To generate yeast expression constructs, the coding sequences of selected ProXs were amplified and inserted into the pDUAL-*Pnmt1*-yeGFP vector^42^. Primers used for vector construction are listed in Supplementary Table 8. The plasmids were linearized with *Not*I restriction enzyme and the resulting fragments were transformed into fission yeast strain LD328 (genetype: *his3-D1 leu1-32*) as described previously^43^. Briefly, yeast cells grown until OD_600_ 0.4-0.8 were collected, washed three times with sterilized water and resuspended in buffer I (240 μL PEG3350 (50% w/v), 36 μL LiAc (1.0 M), 50 μL single-stranded carrier DNA (2.0 mg/mL)). The resuspended cells were then added to 34 μL of the linearized DNA (up to 1 μg), mixed vigorously and incubated at 42°C for 40 min. The cells were resuspended in 100 μL water and plated on EMM+HT (EMM medium supplemented with 45mg/L histidine and 15 μM thiamine). After incubation at 30°C for 2-3 days, individual colonies were selected on EMM+H (EMM medium supplemented with 45mg/L histidine) plates.

### Microscopy

Yeast cells grown on EMM+H plates were inoculated into EMM+H liquid medium and cultured overnight (OD_600_<1.5). 4 μL of cells were placed onto a slide and covered with a coverslip. The imaging was performed on Zeiss LSM880 with Airyscan (Axio Observer 7) Confocal Laser Microscope. GFP was excited at 488 nm and detected at 491-535 nm. Images were acquired with a ×100 objective (NA1.46) and processed by ZEN 3.2 (blue edition).

### Mass spectrometry and data analysis

Samples were separated on 1D SDS-PAGE gel and followed by in-gel digestion, according to the method described before^44^. Briefly, gel bands were excised, discolored, reduced with 25 mM 1,4-dithiothreitol (Inalco, Italy), and alkylated with 55 mM iodoacetamide (Sigma, USA). In-gel digestion was then carried out with trypsin (Promega, USA) in 50 mM ammonium bicarbonate (Solarbio, China) for 16 h at 37°C. The peptides were extracted twice with 0.1% formic acid (Mreda, USA) in 50% acetonitrile (Fisher, USA) aqueous solution. The extracts were centrifuged in a speedvac to reduce volume and were then analyzed by LC-MS/MS.

Peptides were separated by a 120 min gradient elution using a Dionex UltiMate 3000 (Thermo Scientific, USA) HPLC system, which was directly interfaced with a Q Exactive HF-X (Thermo Scientific, USA) mass spectrometer. The analytical column was an Acclaim^™^ PepMap^™^ 100 C18 column (75 μm × 150 mm, 100 Å, 3 μm, Thermo Scientific, USA). The mass spectrometer was operated in data-dependent acquisition mode. For full MS scan, resolution was set to 60,000, scan range was set to 300 to 1800 m/z. For MS/MS scan, a single full MS scan mass spectrum was followed by 40 data-dependent MS/MS scans, resolution was set to 15,000, scan range was set to 200 to 2000 m/z, normalized collision energy was set at 27%.

The MS/MS spectra from each LC-MS/MS run were searched against the proteomes listed in Supplementary Table 6 using the SEQUEST searching engine of Proteome Discoverer (Version 2.1, PD2.1, Thermo Scientific, USA). The search criteria were set as follows: full tryptic specificity was required, 2 missed cleavage was allowed, precursor ion mass tolerance was set to 20 ppm, fragment ion mass tolerance was set to 0.02 Da, oxidation (M) was set as variable modification, carbamidomethylation (C) was set as the fixed modification. The peptide false discovery rate was calculated using Percolator provided by PD2.1. The false discovery rate for protein identification was set to 0.01. Label-free protein quantification was performed by PD2.1. Only unique peptides were used for protein quantification. Missing value was not replaced by minimum value. Quantification result with missing channel was not rejected. Top n peptides used for area calculation was set to 3. Normalization for protein quantification was not employed.

The correlation matrices were generated by GraphPad Prism 9.0.0 software using two-tailed multiple variable analysis. Only proteins that were found in all three biological repeats were used for correlation matrix generation.

## Acknowledgement

We thank Dr. Xiaoyun Zhang for the synthesis of b-isox compound. We thank Prof. Yijun Qi for providing *O. sativa* seedlings, Prof. Junmin Pan for *C. reinhardtii* cells, Dr. Cao Xu for *S. lycopersicum* seedlings, Dr. Tongbing Su for *B. rapa ssp. pekinensis* seedlings, Dr. Haodong Chen for *P. patens* tissues, Prof. Feng Tian for *Z. mays* seedlings and Dr. Zhaorong Hu for *T. aestivum* materials. We thank Dr. Lilin Du for sharing the plasmids and yeast cells used in the heterologous expression system.

## Author contributions

H.Z. and X.F. conceived the study. H.Z. performed b-isox precipitation and confocal imaging experiments and analyses. F.P. performed all bioinformatic analyses. Y.L. performed the mass spectrometry experiments. H.Z., F.P., Y.L., H.D. and X.F. wrote the manuscript, and all authors contributed ideas and reviewed the manuscript.

## Data availability

Full lists of Mass spectrometry were provided as Supplementary Table 1. All the other raw data that support the findings of this study are available from the corresponding authors upon reasonable request.

---

**Extended Data Fig. 1.**
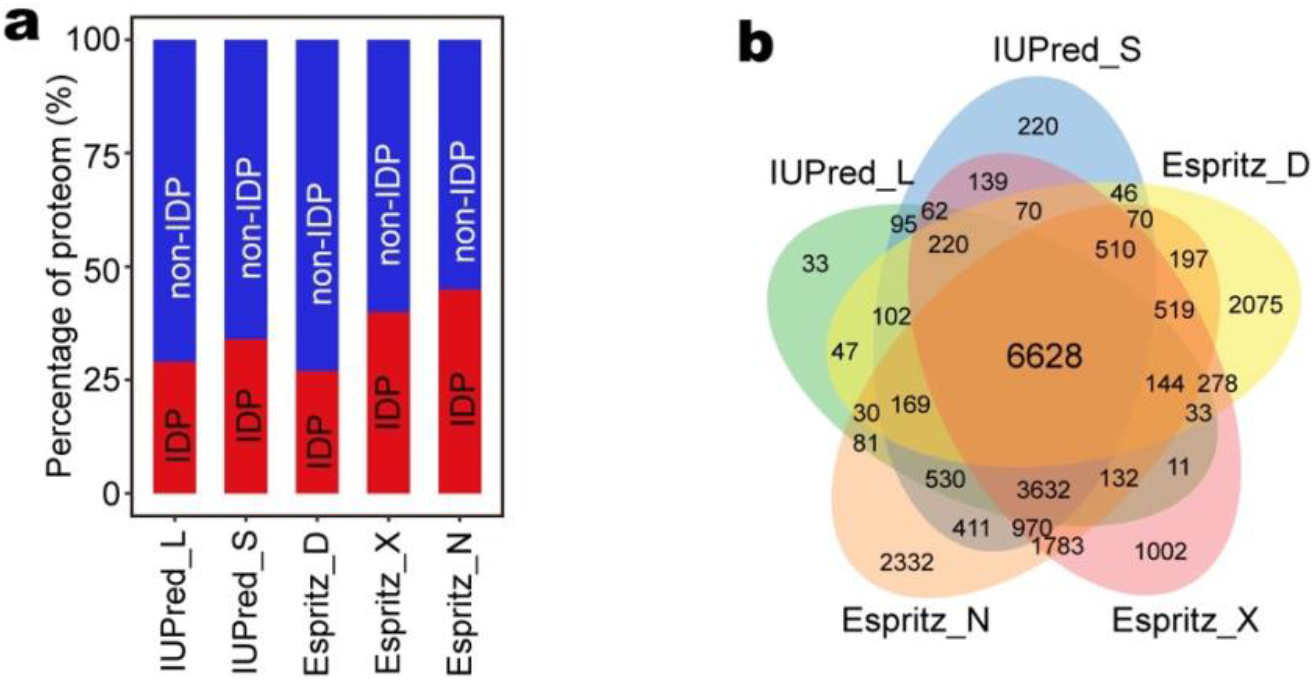
Proteome-wide prediction of IDPs in Arabidopsis. **a**, The percentages of IDPs contained in the proteome of Arabidopsis as predicted by indicated programs. **b**, Venn diagram showing the overlap between the IDPs predicted by different programs.

**Extended Data Fig. 2.**
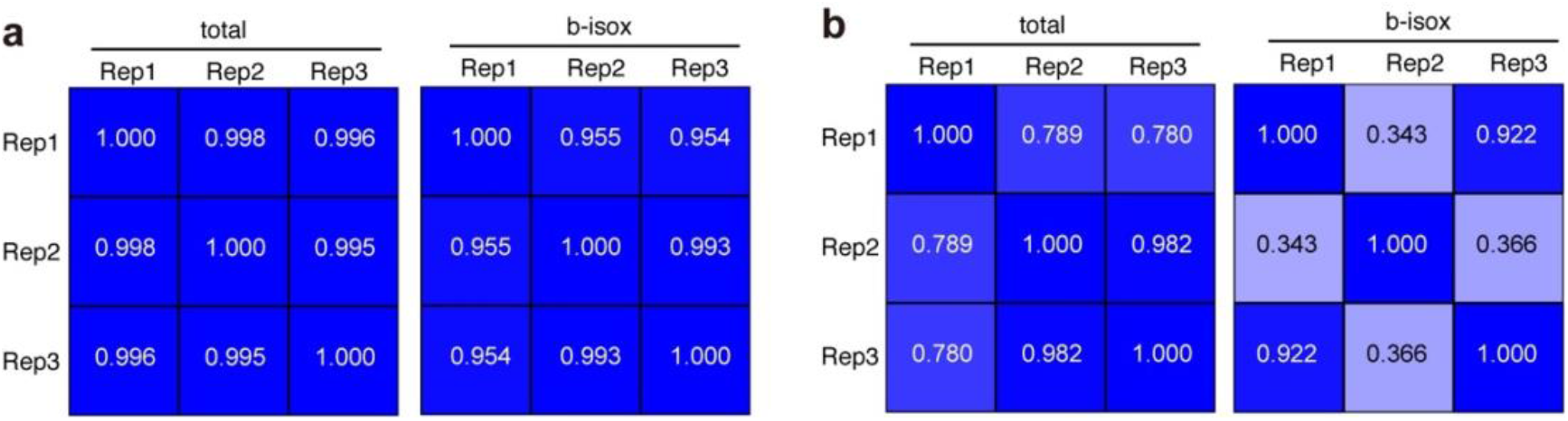
Quality analysis of mass spectrometry data. **a**,**b**, The correlation matrices showing the reproducibility between three biological replicates of indicated protein samples prepared from Arabidopsis seedlings (a) and inflorescences (b).

**Extended Data Fig. 3.**
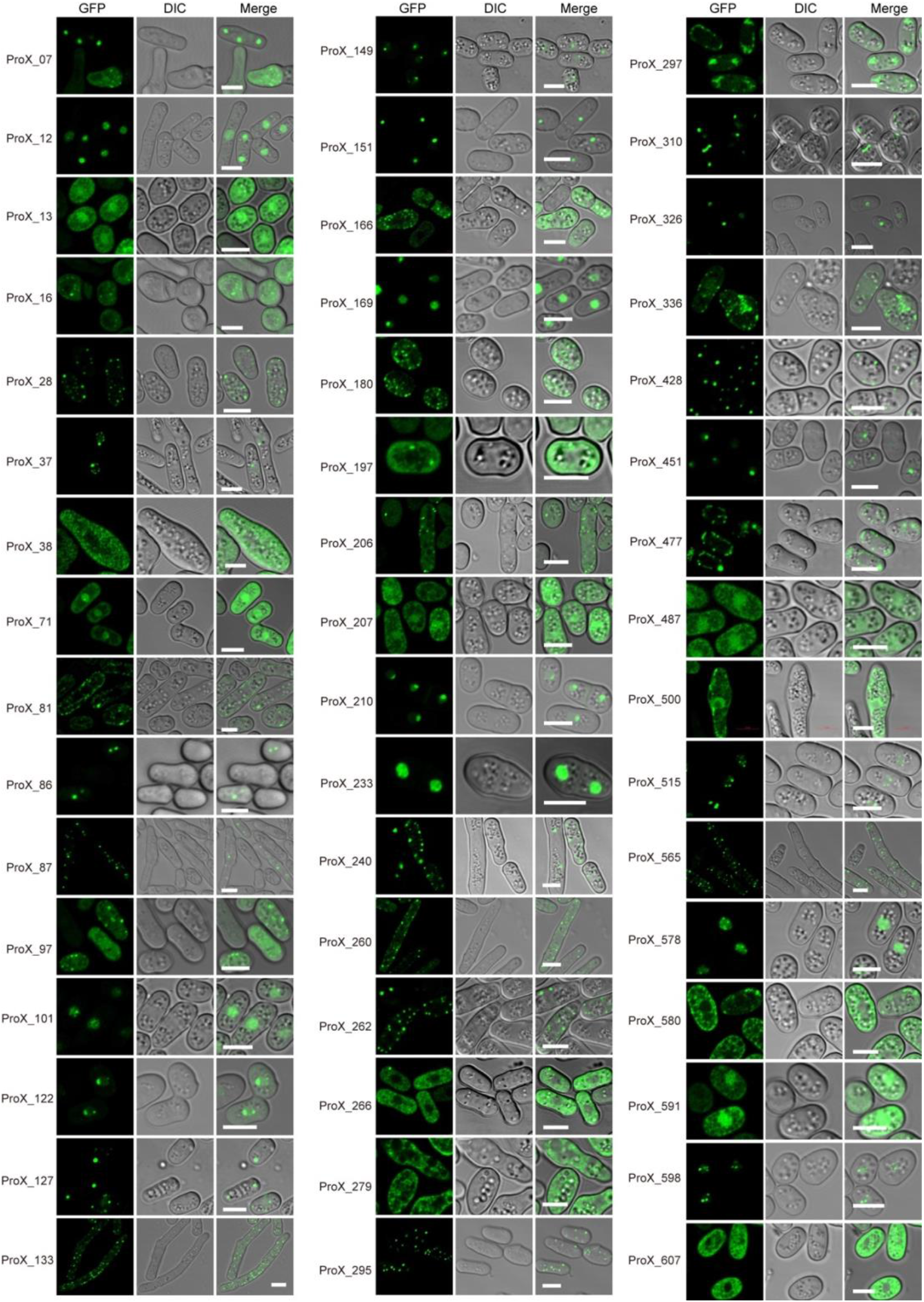
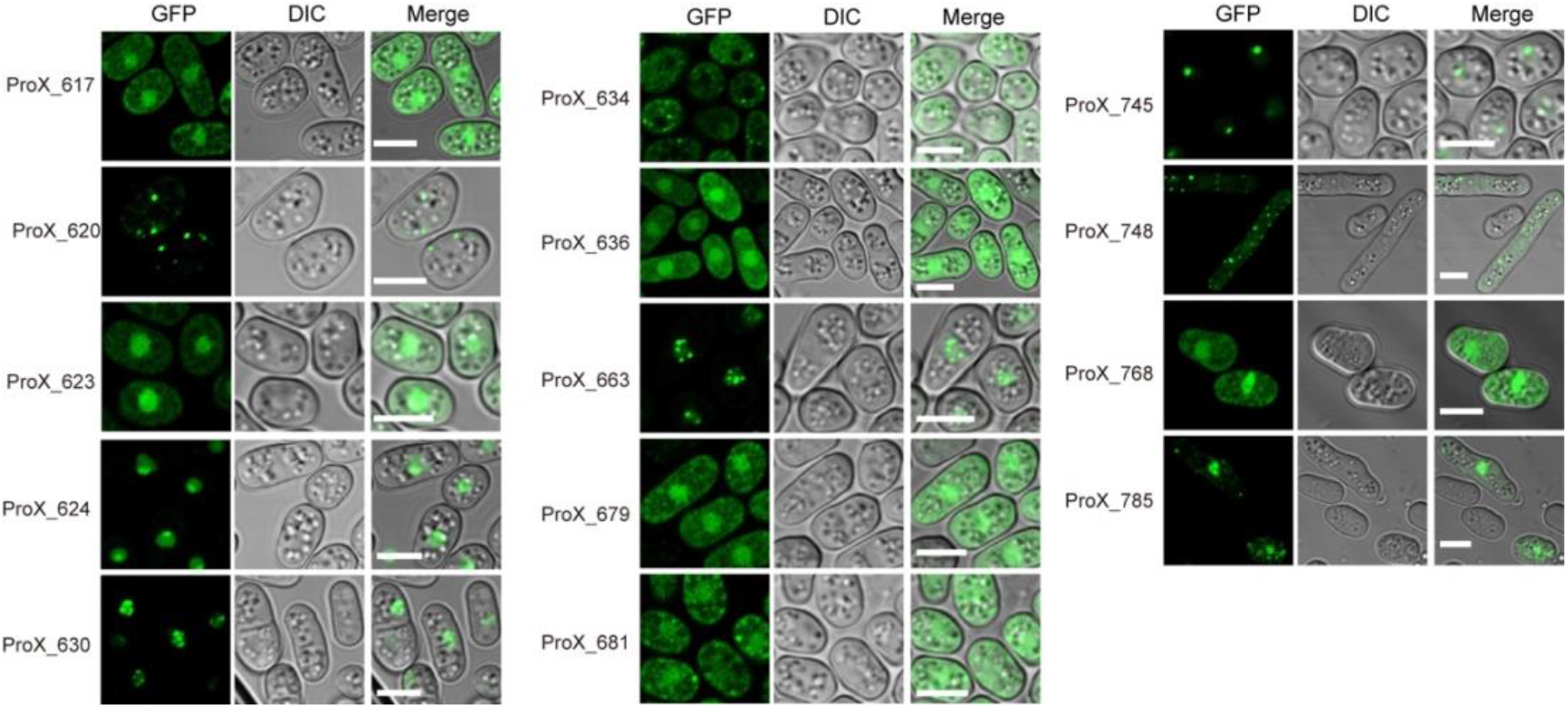
Validation of the ability of ProXs to form membraneless organelles (MLOs). Confocal images of indicated ProXs expressed in yeast cells. Scale bars, 5 μm.

**Extended Data Fig. 4.**
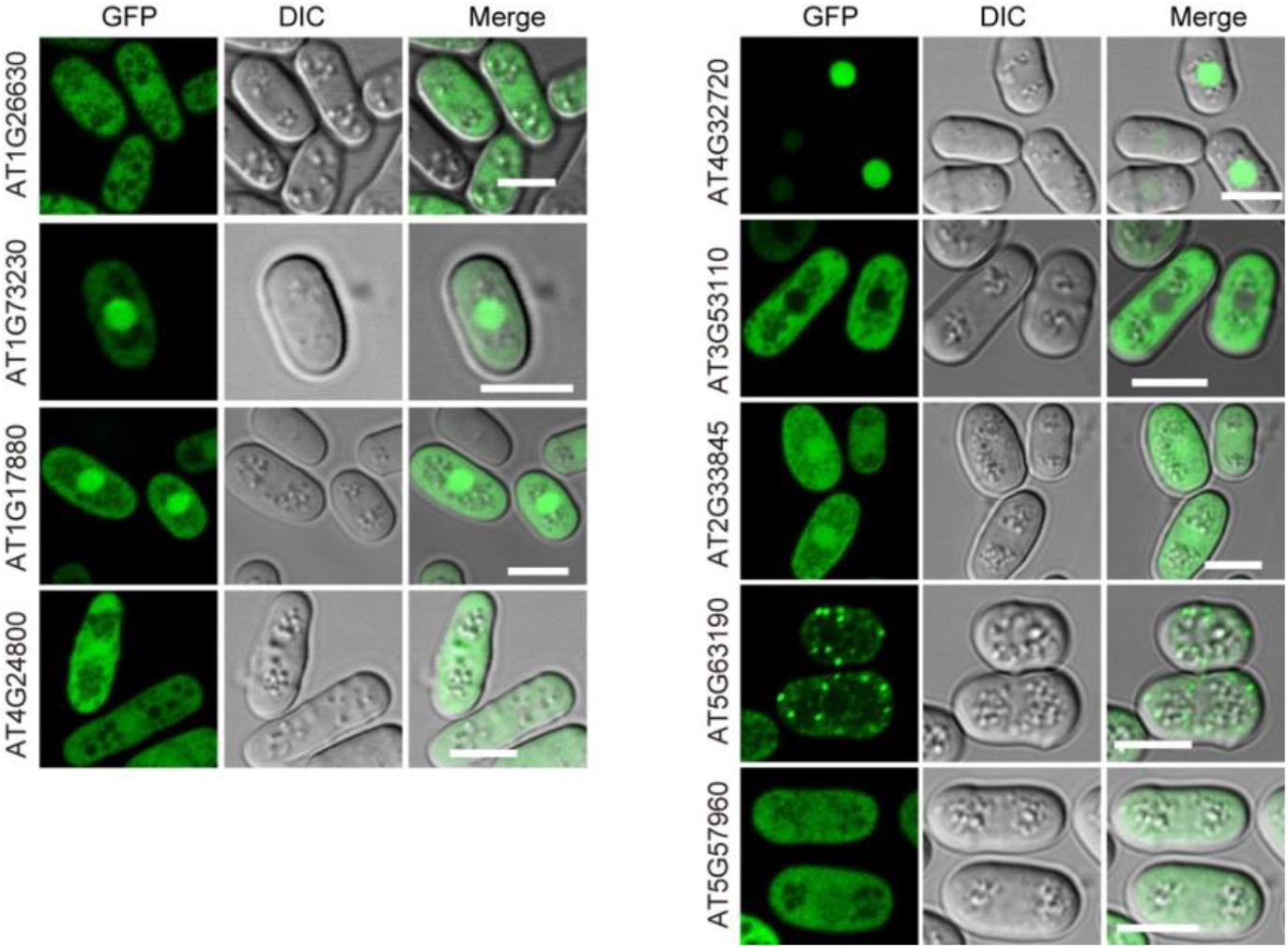
Analysis of RBPs that served as negative controls. Confocal images of indicated RBPs expressed in yeast cells. Scale bars, 5 μm.

**Extended Data Fig. 5.**
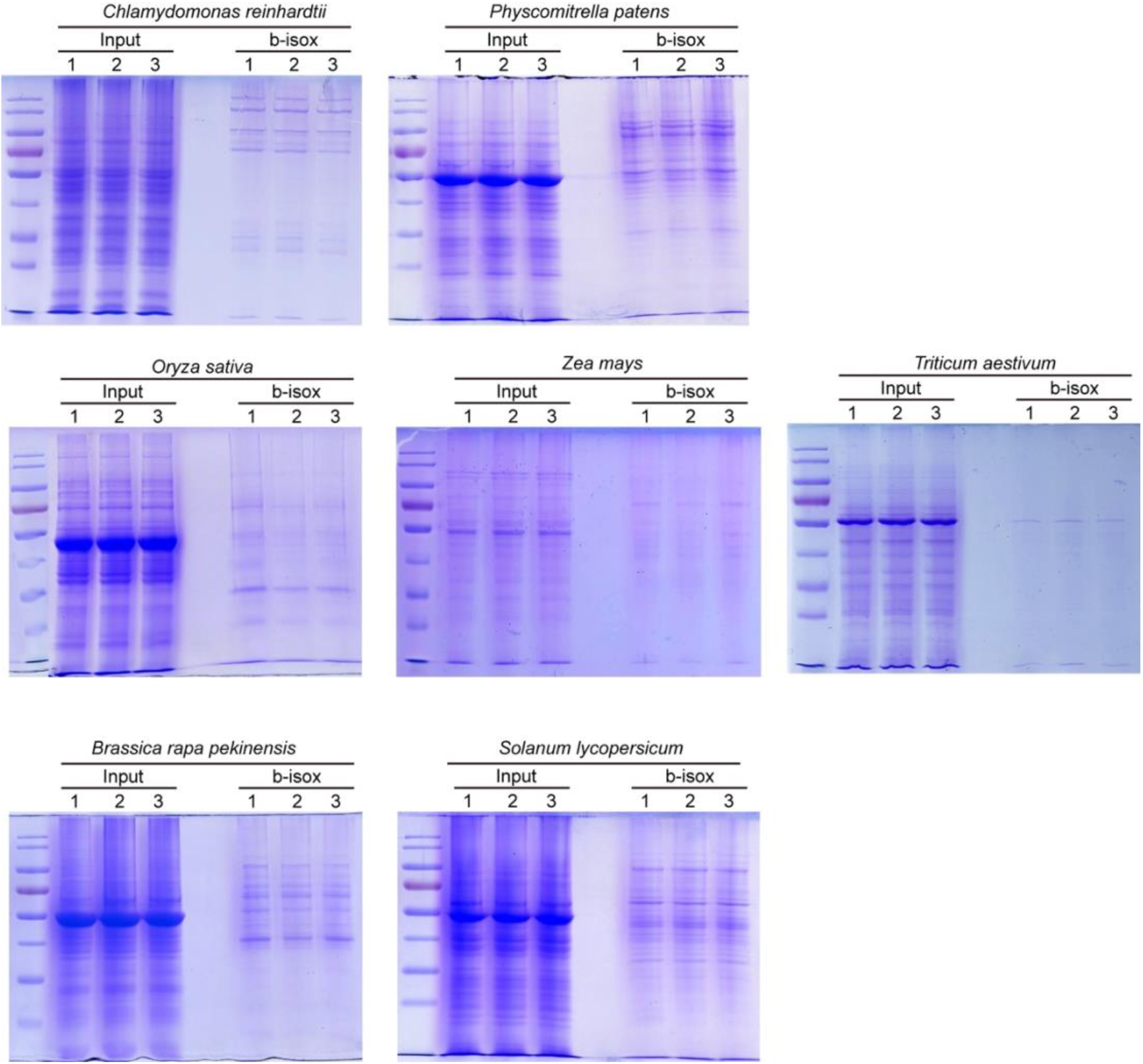
Analysis of protein samples used for mass spectrometry. Coomassie brilliant blue staining of SDS-Polyacrylamide gels separating total and b-isox precipitation samples prepared from the indicated species. The numbers indicate three biological replicates.

**Extended Data Fig. 6.**
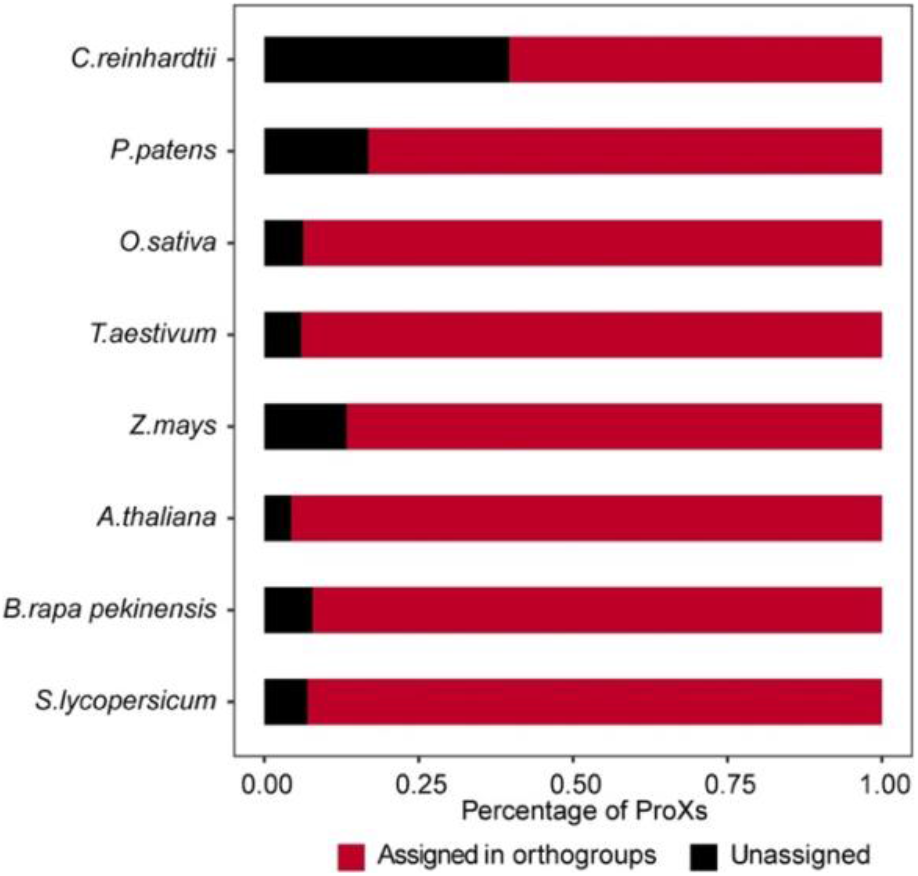
Phylogenetic analysis of the ProXs. The percentages of ProXs that can be assigned into orthogroups in the indicated species.

**Extended Data Fig. 7.**
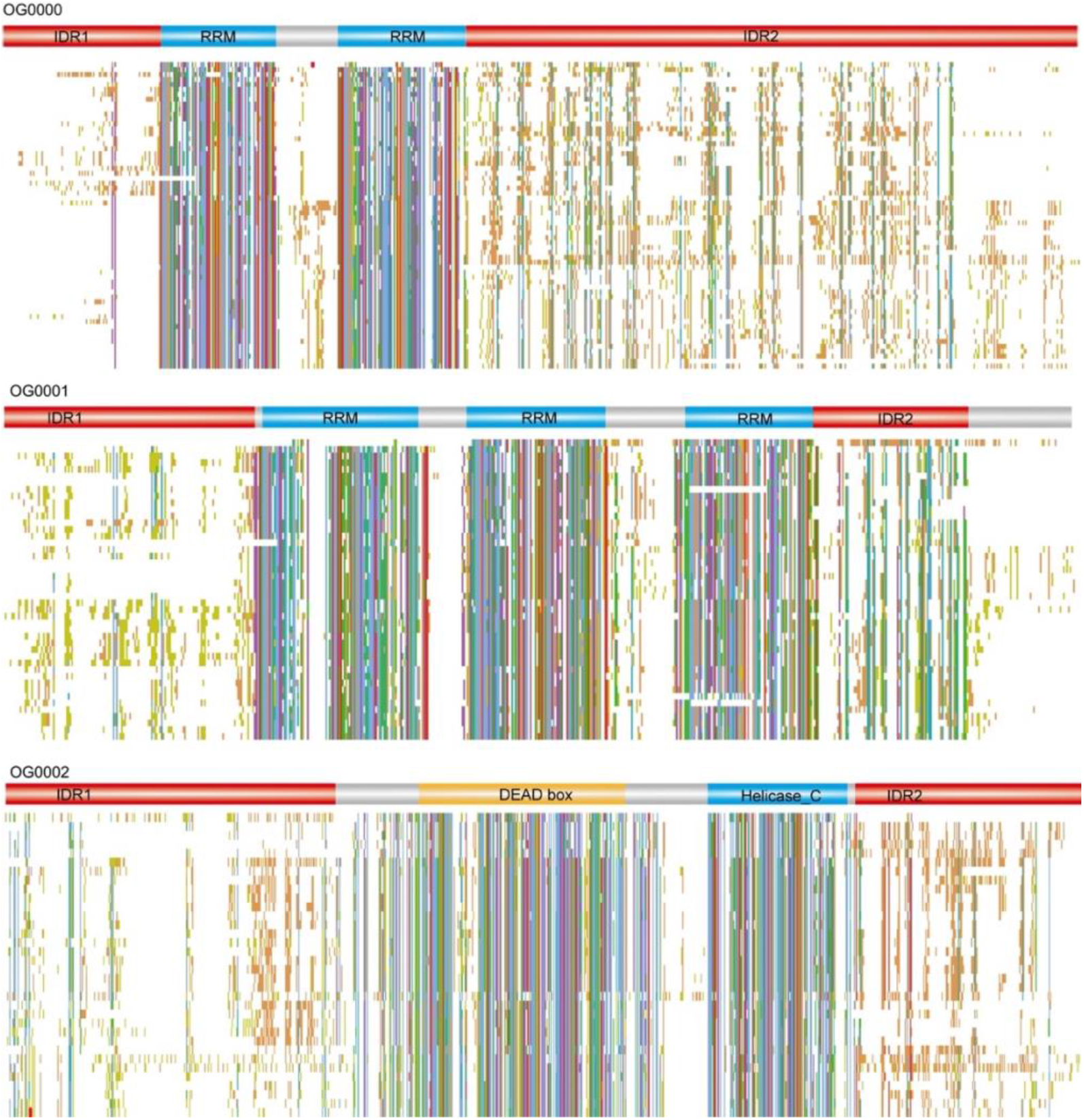
Multisequence alignment of the ProXs reside in the indicated orthogroups.

**Supplementary Table 1. Full lists of ProXs from eight plant species.**

**Supplementary Table 2. A list of AtProXs that were known to form MLOs.**

**Supplementary Table 3. Overlap of AtProXs with RBPomes.**

**Supplementary Table 4. List of AtProXs expressed in yeast cells.**

**Supplementary Table 5. List of control RBPs expressed in yeast cells.**

**Supplementary Table 6. Lists of orthogroups identified in ProXs of indicated species. Related to Fig. 4**

**Supplementary Table 7. A summary of the proteome data collected from each species.**

**Supplementary Table 8. List of primers used in this study.**

